# Mutational interactions define novel cancer subgroups

**DOI:** 10.1101/187260

**Authors:** Jack Kuipers, Thomas Thurnherr, Giusi Moffa, Polina Suter, Jonas Behr, Ryan Goosen, Gerhard Christofori, Niko Beerenwinkel

## Abstract

Large-scale genomic data can help to uncover the complexity and diversity of the molecular changes that drive cancer progression. Statistical analysis of cancer data from different tissues of origin highlights differences and similarities which can guide drug repositioning as well as the design of targeted and precise treatments. Here, we developed an improved Bayesian network model for tumour mutational profiles and applied it to 8,198 patient samples across 22 cancer types from TCGA. For each cancer type, we identified the interactions between mutated genes, capturing signatures beyond mere mutational frequencies. When comparing mutation networks, we found genes which interact both within and across cancer types. To detach cancer classification from the tissue type we performed de novo clustering of the pancancer mutational profiles based on the Bayesian network models. We found 22 novel clusters which significantly improved survival prediction beyond clinical and histopathological information. The models highlight key gene interactions for each cluster that can be used for genomic stratification in clinical trials and for identifying drug targets within strata.

## Introduction

The past years have seen great progress towards a deeper understanding of the molecular changes underpinning cancer progression. Identification and characterisation of molecular subtypes within and across different cancer types has emerged as a promising approach for the development of targeted therapies [Sun et al., 2007, Higgins and Baselga, 2011, Roock et al., 2011]. Nevertheless, cancer treatment is far from optimal. The approval of the limited number of available cancer drugs is often limited to a specific cancer type or subtype, preventing widespread use of targeted therapies. Moreover, cancers may develop resistance against these therapies, rendering them ineffective [Groenendijk and Bernards, 2014].

Pancancer analyses enabled by large datasets such as The Cancer Genome Atlas (TCGA) [McLendon et al., 2008] or the International Cancer Genome Consortium (ICGC) [Hudson et al., 2010] may aid a better understanding of the disease biology across different tissues, and can, for example, identify mutational hotspots in tumours [Chang et al., 2016]. Genes strongly associated with a cancer type or novel pancancer subgroups provide insights into the molecular mechanisms, which are key for pinpointing novel therapeutic opportunities and improving current treatment strategies. For example, analysis of endometrial carcinoma showed genetic similarities to certain types of breast and ovarian cancer [TCGA Research Network, 2013] while olaparib, approved for BRCA-mutated ovarian cancer, provided a good response in metastatic prostate cancer patients with DNA repair mutations [Mateo et al., 2015]. Pancancer, or basket, clinical trials [Cunanan et al., 2017] could extend targeted treatments beyond their current indication, like testing the *BRAF* inhibitor vemurafenib outside of metastatic melanoma [Hyman et al., 2015], or test novel agents, like Loxo Oncology’s trial of larotrectinib for patients with a *TRK* gene fusion mutation. Recently, the FDA granted its first approval for a drug (pembrolizumab) based on genetic markers, regardless of the tissue type.

Cancer is known to be a disease characterised by a progression of molecular changes leading to malignant features and activities [Nowell, 1976, Vogelstein et al., 1988]. Alongside stratifying tumours based on static molecular profiles [Ciriello et al., 2013, Hoadley et al., 2014], investigating their development [Gerstung et al., 2009, Attolini et al., 2010, Gerstung et al., 2011, Farahani and Lagergren, 2013, Misra et al., 2014, Ramazzotti et al., 2015, Cristea et al., 2017] may offer a new perspective on pancancer analyses with the potential to identify key drivers and provide benefits on multiple levels, including (1) prioritisation of mutation-based biomarkers; (2) uncovering previously unknown mutational dependencies; (3) identification of biomarkers of progression; and (4) biological insight into the genetic progression of cancer.

The facets of clustering patient samples, inferring their genetic tumour progression and mutational interactions are highly inter-related (Supplementary Material A). Here, we introduce a unified statistical framework to combine them by modelling the mutations as a Bayesian network. The probability of observing each mutation depends on the state of its parents in the network, thereby accounting for mutational interactions such as co-occurrence or mutual exclusivity, as well as more complex relationships. The directions of the connections may be suggestive of causal relationships [Pearl and Verma, 1991, Pearl, 2000, Dawid, 2010], though they may not be fully resolved from the data.

We developed novel and efficient methods to infer the dependency structure of the mutations and performed fully Bayesian inference (Methods) to capture the uncertainty in the network structure learned from mutational profile data. Characterising mutational data through Bayesian networks can provide novel and useful insights through the analysis of the mutational interactions encoded by the network, beyond just analysing mutational frequencies. We employ our Bayesian network modelling to cluster patient samples into groups, with different interactions among mutated genes. The key interactions within and across novel subgroups may uncover common mechanistic insights, potential therapeutic targets, and prognostic and predictive biomarkers.

## Results

We performed two distinct analyses of nonsilent mutation data, summarised at the gene level for 201 genes, from 8,198 patient samples across 22 cancer types (Supplementary Material Table S1) from TCGA. Initially, in a supervised analysis, we built cancer-specific probabilistic models to explore type-specific mutational interactions and pancancer heterogeneity. Then, in an unsupervised analysis, we proceeded to cluster the samples into novel mutational subgroups (Figure 1).

**Figure 1:**
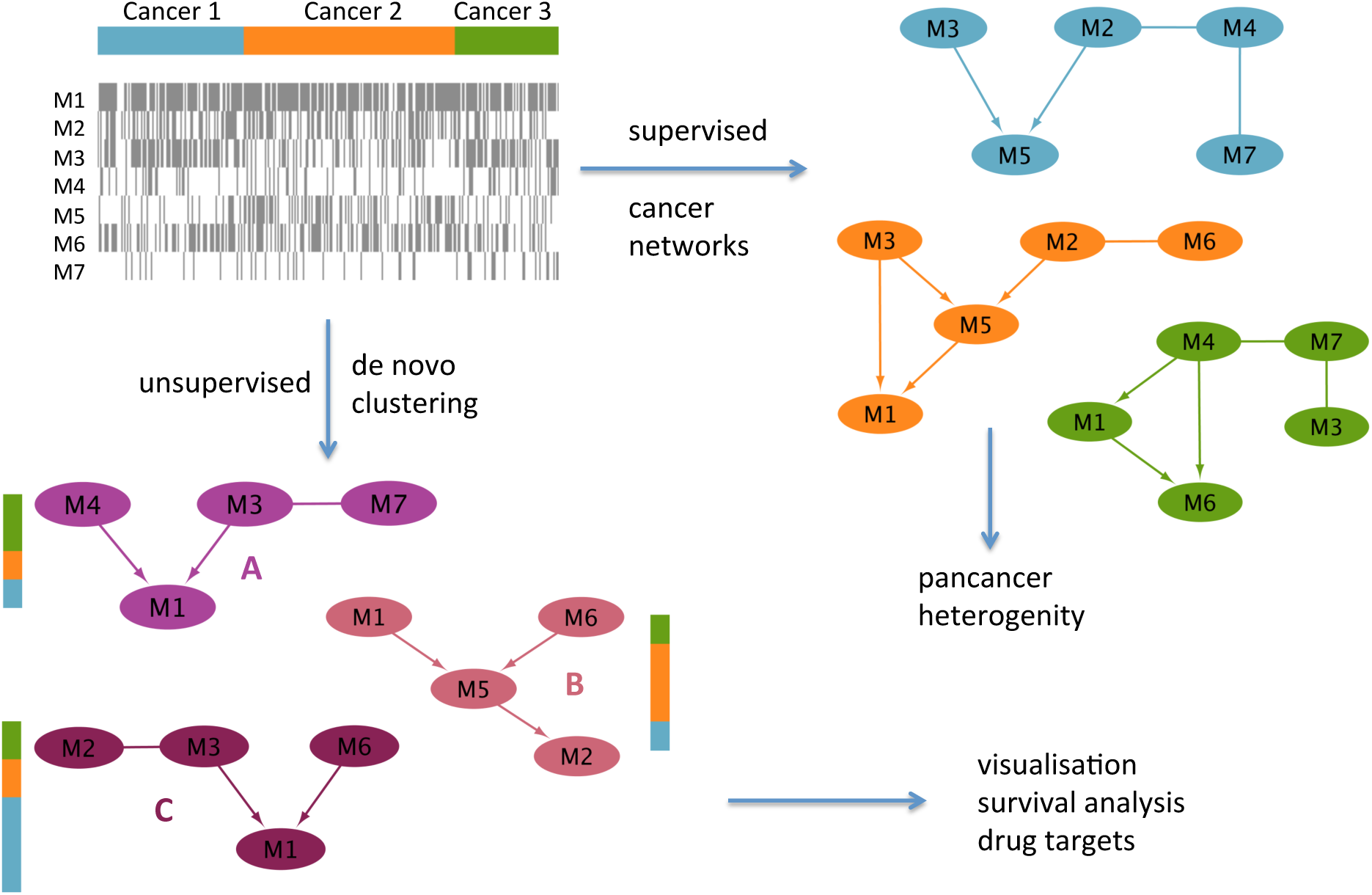
Starting from the mutation data, we perform two types of analysis. Supervised learning of the Bayesian network structure for each known cancer type, allowing us to uncover mutational interactions and visualise pancancer heterogeneity. Unsupervised clustering of the mutation data into components with common interactions to uncover a novel stratification of the patient samples.

### Cancer-specific Bayesian networks

Stratifying the TCGA mutation data by tissue of origin, we built Bayesian networks (Methods) separately for to each cancer type. We obtained an alternative representation of the mutational landscape which goes beyond mutation frequencies by highlighting the interdependencies between genes. Edges show co-occurrence or mutual exclusivity, or higher order correlations between sets of genes, offering a systematic visualisation of the most important mutational interactions in 22 different cancer types (Figure 2).

**Figure 2:**
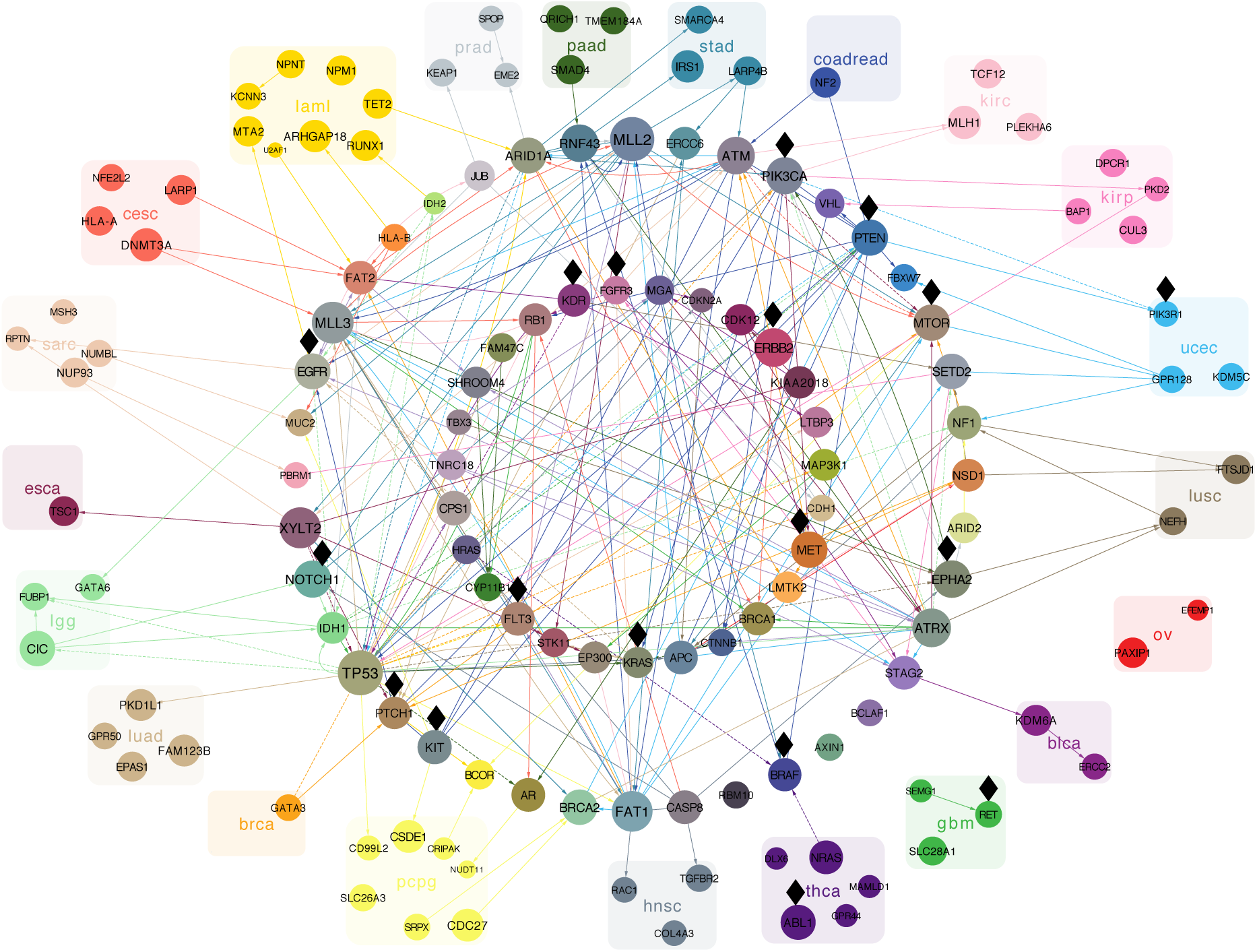
The cancer-specific connections between the genes, limited for clarity to only display connections between the 20 most frequent and connected mutations per cancer type. The Bayesian networks for each cancer type are overlaid on the same union of gene nodes. Edges highlight interactions between the selected mutations and are coloured by the cancer type for which they occur (leukaemia: yellow; glioblastoma: green, etc.). Directed edges point in the inferred direction of dependency, whereas undirected edges indicate cases where the direction cannot be inferred. Solid edges indicate positive correlation (a degree of co-occurence) while dashed edges indicate negative correlation (a degree of exclusivity) between the genes connected. Genes are coloured according to the colours of their edges. Those with edges in only one specific cancer type are grouped and labelled with the same colour. Genes with edges in different cancer types are arranged in the two circles in the centre. The size of each gene correlates with the total number of edges across all cancer types, including edges that are not shown. Black diamonds mark putatively actionable targets.

The network parameters estimated from the data capture information both about the mutational frequencies as well as their interactions, allowing us to evaluate how well each model explains the mutational status of each patient sample (Methods), including those from other cancer type. Simulations show that our approach performs notably better than alternatives in learning the network structure (Supplementary Material B). It also improves on the potential to effectively characterise different cancer types with respect to simple distance-based measures (Supplementary Material C). The network structure inference is informed (Methods) by using the STRING protein-protein interaction network [Szklarczyk et al., 2014] as a prior, but even without this information a comparison (Methods) reveals a significant overlap of the edges (permutation test; *p* = 4.1 × 10^−5^), suggesting the inferred network of mutational interactions is biologically relevant as known functional interactions coincide.

We rediscovered many key genes which have previously been associated with their respective cancer types. For example, genes *ATM, PIK3CA* and *PTEN* with a lot of connections in colorectal cancer (Figure 2) have previously been reported as highly mutated [TCGA Network, 2012]. In lung adenocarcinomas, we recapitulate the mutual exclusivity between *KRAS* and *EGFR* as well as the importance of *TP53* [TCGA Research Network, 2014], for which our model suggests that it is frequently co-mutated with both *KRAS* and others like *MLL3*, as well as *KRAS* and *STK11* are frequently co-mutated.

*TP53* is a major hub with the most interactions across multiple cancer types (67 in total) and with multiple dependencies in brain cancer and especially lower-grade glioma. *TP53* mutations in lower-grade glioma have previously been associated with the disease, along with *IDH1, FUBP1, ATRX, CIC, NOTCH1, EGFR*, and *PIK3CA* [TCGA Research Network, 2015] amongst which we see some interactions. Similarly in glioblastoma, where in addition to *TP53*, we observe interactions involving *PTEN, IDH1* and *ATRX* which were previously reported to be mutated [Brennan et al., 2013].

Other hubs with high connectivity like *TP53* include *MLL2* (65 interactions), *MLL3* (58), *XYLT2* (56) and *FAT1* (55). Mutations in these genes have previously been associated with cancer [Olivier et al., 2009, Morris et al., 2013, Kantidakis et al., 2016]. *MLL3* and *FAT1* share connections across several cancer types, with both having several interaction in uterine cancer, while *MLL2* exhibits many connections for stomach, and *XYLT2* for esophageal cancer.

Strong edges can hint at common mechanistic causes or fitness effects. For the example of *FAT1*, our study finds relevant interacting mutations in breast, colorectal, endometrial, kidney, lung, liver, stomach and head and neck cancer, with mutation rates between 2% – 24%. Although, *FAT1* was reported to be recurrently mutated in several cancer types [Katoh, 2012, Garg et al., 2015, Morris et al., 2013], our results suggest that the gene correlates with a large number of mutations in different cancer types, including *FAT1* in breast cancer, *ATM* in colorectal cancer, and *APC*, *MTOR* and *MLL3* in endometrial cancer. On the pathway level, we found highly connected genes across cancer types to be significantly involved in many signal transduction pathways, as well as cellular processes and DNA damage repair (Supplementary Material Table S8).

Several genes with mutational interactions are putatively actionable, with drugs either approved or currently tested in a clinical study (labeled with black diamonds in Figure 2). In general, approval of targeted therapies is limited to one or several cancer types or cancer subtypes. Strong dependencies in the Bayesian network potentially indicate effectiveness of targeted therapies against dependent gene mutations, particularly if it is in the same pathway. Moreover, the network potentially expands the group of tumours responsive to a specific targeted therapy, within the same tumour types and between different tumour types. Although the drugs are only targeting the mutation itself, we see for example several interactions of *KRAS* in lung adenocarcinoma (with *TP53, EGRF* and *STK11)* and there are currently several clinical trials investigating the effectiveness and safety of targeted therapies against *KRAS* for this cancer type (NCT02642042, NCT01912625 or NCT02079740).

A two-dimensional visualisation based on multiscale projections (Methods) highlights the differences and similarities of patient samples as measured over the Bayesian networks, within and across cancer types (Figure 3). Some, like colorectal, thyroid and lower grade glioma, are well separated and hence well defined by the mutation profiles of their patient samples; others are much more similar, for example those within overlapping groups. Even for the better separated cancer types, we observe substantial heterogeneity with some patient samples closer to those from other cancers.

**Figure 3:**
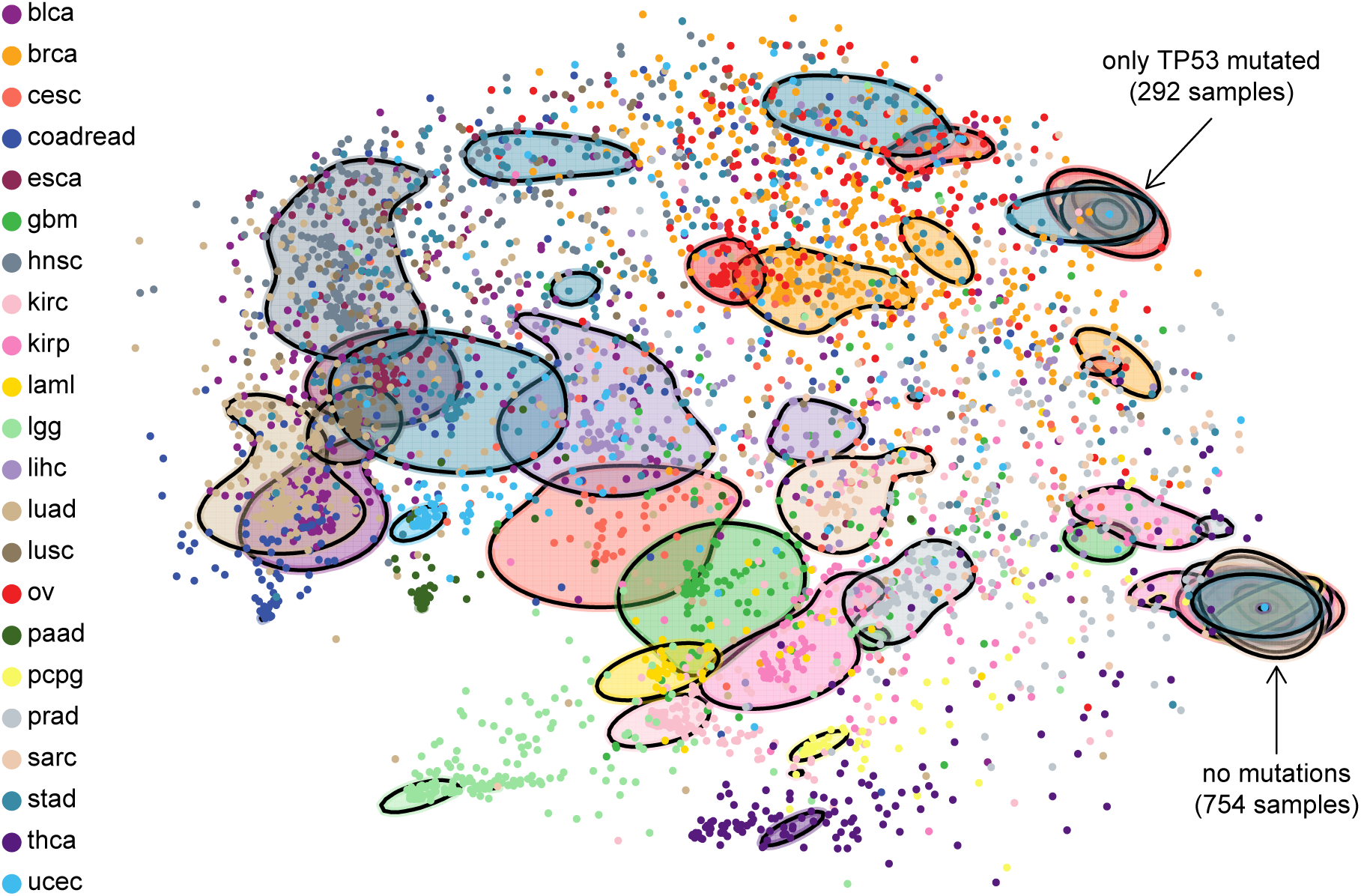
2D visualisation of the similarity between patient samples based on their fit to each cancer-specific Bayesian network, highlighting the heterogeneity within and across cancer types. For example, stomach, breast and liver cancer show high inter-tumour heterogeneity with a high spread across the plot, whereas pancreatic cancer shows low inter-tumour heterogeneity and is much more localised. Ovarian and breast cancers as well as bladder cancer and lung adenocarcinomas show similar mutational profiles, while lower grade glioma is rather distinct from other cancer types, as is thyroid cancer. The solid shapes are based on contours which together contain a total of 50% of the respective cancer types. The group of samples on the lower right possess no mutations among the 201 genes while those exhibiting a mutation only in *TP53* are also indicated. Versions highlighting certain cancer types are displayed in Supplementary Material Figure S5.

As well as being applicable to pancancer cohorts, these methods naturally apply when focussing on individual cancer types and their subtypes (as we discuss for breast cancer in Supplementary Material D).

### De novo cancer type-independent clustering based on Bayesian networks

Given the considerable heterogeneity across cancer types, we asked ourselves whether the mutation profiles themselves can be re-clustered on the basis of Bayesian network models without knowing the cancer types. This model-based de novo clustering of the binary mutation data (Methods and Supplementary Material F) identified 22 groups, coincidentally the same number as the original cancer types (which was not imposed), but distinct in their composition. Each cluster is defined by a Bayesian network which constitutes a generative model of its patient data. These models fit the data much better than the partitioning by cancer type – the average cluster assignment is 91.5% compared to 71.9% for the cancer specific models.

The composition of the clusters (Figure 4) shows that the more differentiated cancer types, like colorectal, thyroid and lower grade glioma from the 2D projection (Figure 3) initiate new clusters. Clusters G and I are mostly composed of glioma and colorectal samples respectively. Almost all thyroid cancer samples belong to cluster V, along with samples from other cancer types, and this cluster exhibits a strong enrichment in *BRAF* mutations (17% compared to an overall rate of 7%, Supplementary Material Table S9). Cluster K, where *TP53* plays a strong role, mostly consists of colorectal samples, and cluster L of leukaemia with *IDH* mutations being prominent. The ovarian cancer samples belong almost entirely to cluster U (where all samples exhibit a *TP53* mutation). Samples with no mutations among the 201 genes are assigned to cluster V.

**Figure 4:**
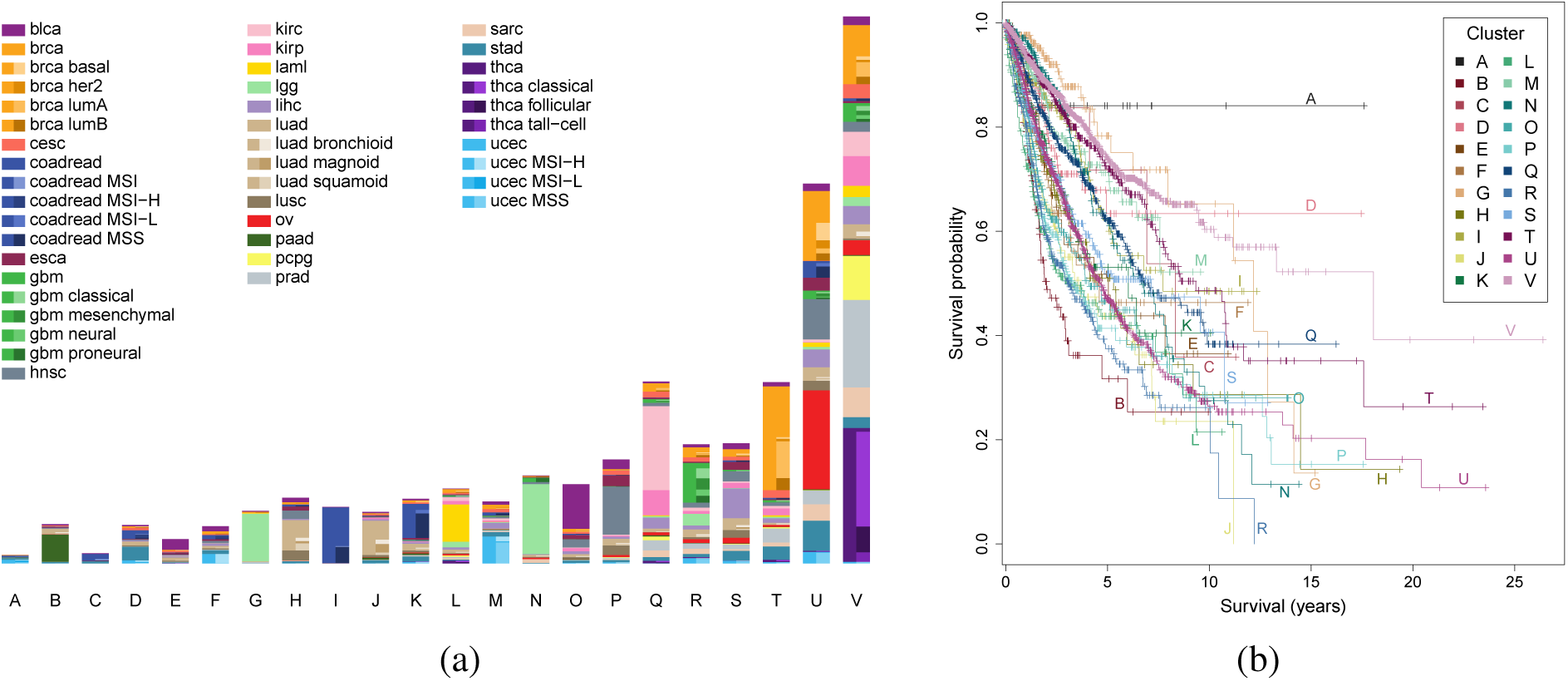
(a) Assignment of the 8,198 patient samples to the 22 clusters, labelled A through V, based on Bayesian network clustering of their mutation profiles. The left hand side of the bar for each cluster indicates the number of patient samples with a given cancer type, while the right hand side indicates the breakdown into the known subtypes. (b) Survival probabilities of the 22 clusters.

The de novo clustering also split patient samples from the same cancer type, like the glioma samples into clusters G and N: cluster N has elevated mutation rates in *TP53* and *ATRX* which are largely absent from cluster G which instead has elevated rates of *CIC* as well as even higher rates of *IDH1* than N (98% compared to 86%). Patient samples from certain cancer types may actually be more similar to tumours in other cancers. Cluster S, for example, groups samples of several different cancer types, including bladder, breast, liver and lung cancers and possesses elevated mutation rates in several genes, like *TP53, MLL, ARID* and *CTNNB1*.

For cancers with known subtypes, the clustering recovered some of these differences. For example, the microsatellite stable subtype of uterine cancer has a rather different cluster composition than the other uterine patient samples (Supplementary Material Figure S14), with enrichment moving from clusters M and U to A and F. Similarly, the colorectal subtypes are quite distinct in their cluster composition. For the breast cancer patient samples, the luminal subtypes have a strong presence in clusters T and V while the basal and Her2 enriched subtypes are mostly absent from these clusters and appear preferentially in cluster U instead.

The clusters are highly significant in predicting survival (likelihood ratio = 38.3; *p* = 2.9 × 10^−8^), also accounting for clinical and histopathological information (age, stage and cancer type; Methods), showing that considering the mutation data and their interactions provides a more complete and informative picture of tumours and their progression. The Bayesian network clustering also performs favourably in survival prediction compared to a range of standard clustering approaches (Supplementary Material Table S6).

The clustering is based solely on the mutation data, so we also compared individual clusters without adjusting for clinical and histopathological information. We observed significant differences (Figure 4 and Table S3) showing that the clustering uncovers strong biological signals. In particular clusters G, M, T and V show good survival (five-year overall survival of 70 — 80%), whereas clusters B, P and R show poor survival; cluster B has the poorest outcome with a five-year overall survival of just over 30%.

Some of the differences between individual clusters can be explained by the mutational profiles correlating with and recapitulating clinical and histopathological variables. When we remove the confounding by age, stage and cancer type, clusters G, J, N and U are still significantly different compared to reference (cluster V, Supplementary Material Table S4). Clusters G and N both contain a large fraction of lower grade glioma patient samples and have good survival. The cancer type is adjusted for in the survival analysis, meaning that the improvement in survival predictions is due to the diverse patient samples that are clustered with them (Figure 4) along with the separation of lower grade glioma into two main groups. Clusters J, with a large group of lung adenocarcinomas, and U have significantly poorer prognosis.

Some cancer types are split by the clustering, with one portion forming the bulk of a cluster, and show differences in survival between the patients assigned to different clusters. For example, colorectal samples in cluster K show a significantly higher lethal risk than colorectal samples in cluster I (hazard ratio = 1.7; *p* = 0.04). This is in line with the fact that cluster K contains a large number of samples from the MSS subtype, which has the worst outcome [Phipps et al., 2015]. Similarly, leukaemia samples in cluster L show a significantly lower lethal risk compared to samples in cluster V (hazard ratio = 0.5; *p* = 0.01).

The probabilistic model describing each cluster can be utilised to visualise the important mutational interactions characterising the mutation profiles of its samples (Figure 5). The cluster composition is distinct from the grouping by cancer (Figure 2), even for clusters dominated by one cancer type such as cluster B, G, I or J, so that the key interactions mostly differ, especially when focussing on 20 genes per cluster for clarity.

**Figure 5:**
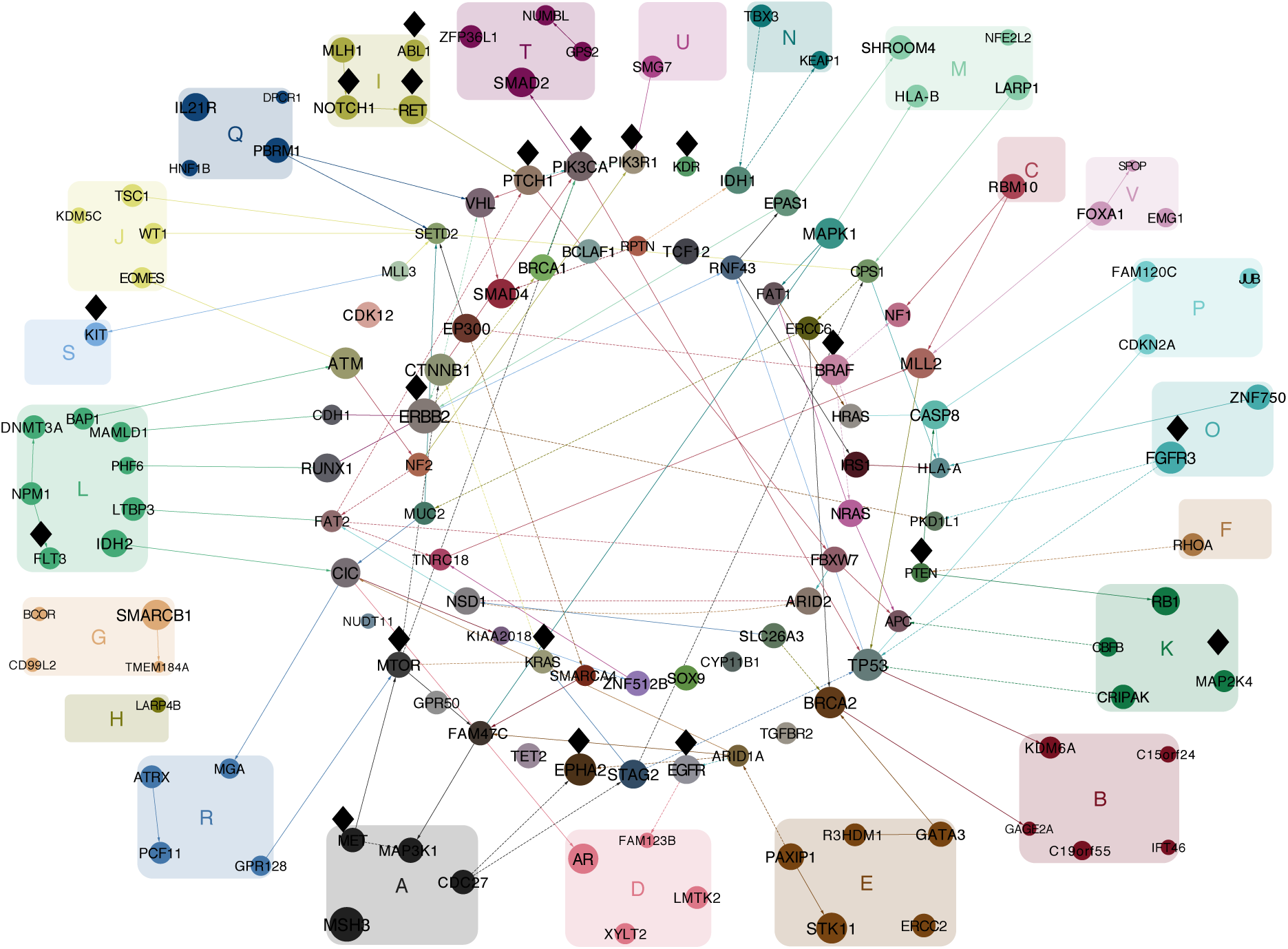
The connections between the 20 most frequent and connected genes per cluster (rather than per cancer type, as in F**********igure 2). Black diamonds near nodes indicate putatively actionable genes. Edges highlight interdependencies between the selected mutations: solid for positive and dashed for negative correlation. Directed edges point in the inferred direction of dependency, whereas undirected edges are where the direction cannot be inferred. Node size reflects the total number of edges, including edges not shown. Nodes are coloured by combining the colours of their edges from the different clusters.

*TP53*, as the most frequently mutated gene across all cancer types, remains fairly prominent when clustering samples by their mutation profile. Looking at its connections among all genes, these differ substantially across the clusters even when the marginal frequency is similar (Figure 6), suggesting that it plays different roles in different clusters. Our approach emphasises mutational interaction patterns which are specific to a cluster. Along with its interactions, the prevalence of *TP53* in each cluster is also used in assigning patient samples. For example all members of cluster U possess a *TP53* mutations while none of cluster V do.

**Figure 6:**
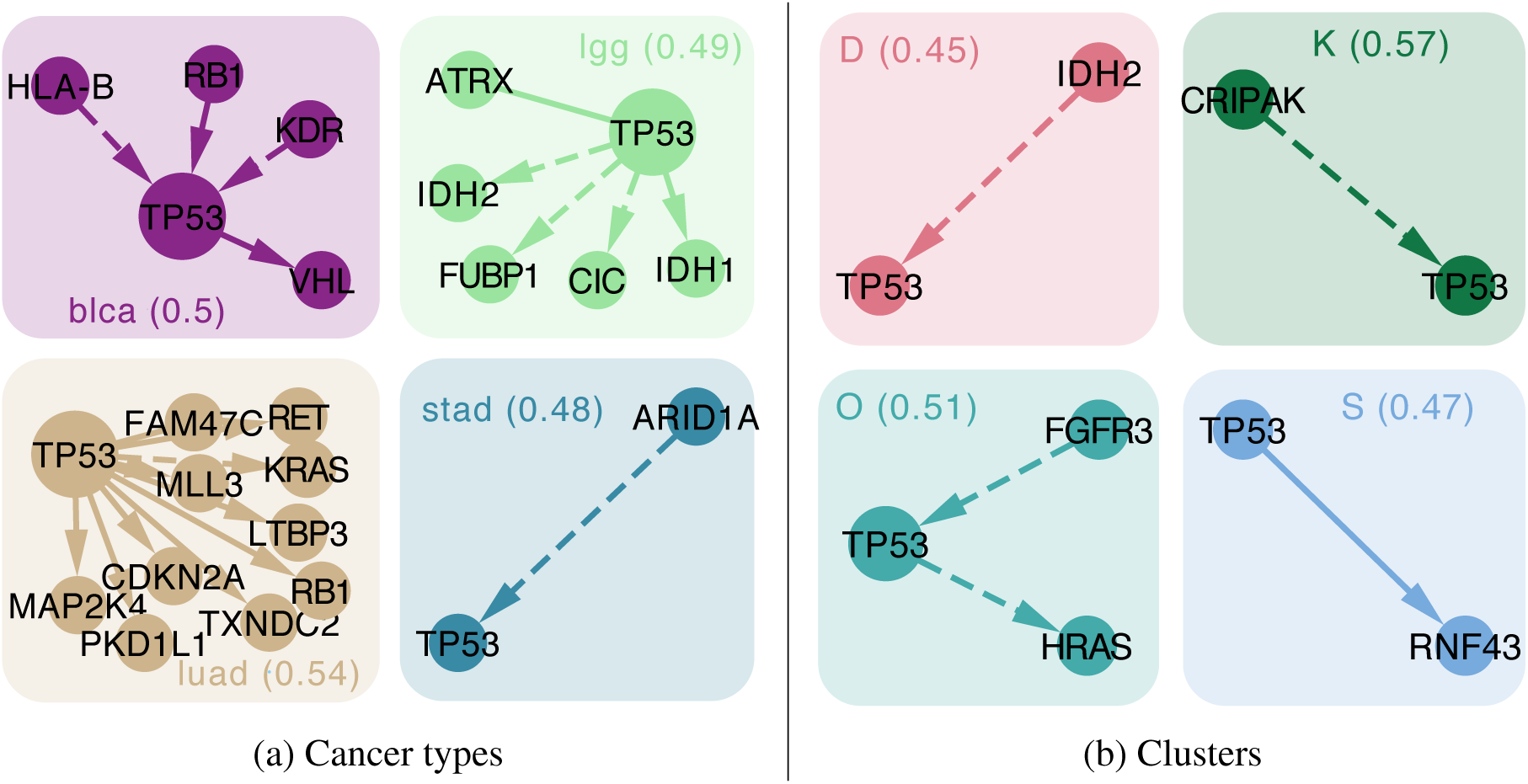
Focus on *TP53:* (a) Cancer types or (b) clusters are characterised by interactions of *TP53* with different genes. For cancer types or clusters where *TP53* shows a similarly high mutation frequency (around 0.5), we display all its interactions (positive correlation is represented by solid arrows, negative correlation by dashed). Groups are annotated by the cancer type or cluster name and the respective mutation frequency of *TP53* across samples (in brackets). Several of these interactions have been studied in cancer. For example, an inverse relationship between *ARID1A* and *TP53* was previously observed in gastro-intestinal cancers [Wu et al., 2014] while *TP53* mutations co-occur with *RB1* in bladder cancer [Longo et al., 2016].

Among the clusters, genes such as *ERBB2* (37 interactions), *MSH3* (36) or *CDKN1B* (33) have more interactions than *TP53* (32) in total. Of the selected interactions (Figure 5), *ERBB2* has connections particularly for clusters T and M, and for cluster M interacts with *CTNNB1. PIK3CA* has several interactions, including three for cluster C (with *TP53, CTNNB1* and *VHL)*, a cluster dominated by colorectal cancer samples.

We found several cancer-associated signalling pathways significantly enriched in highly connected genes across clusters, including HIF-1, Jak-STAT, p53, Toll-like receptor, and TNF signalling (Supplementary Material Table S10). These results suggest that genes involved in signalling pathways are not only highly connected, with a large number of mutational interactions with other genes, but that they also play an important role in characterising the clusters.

Clustering samples by their mutational profile, independent of their cancer type, provides an alternative patient stratification which may inform targeted treatment (putatively actionable genes are marked by black diamonds in Figure 5). For example, *BRAF* inhibitors are being currently tested or have already been approved for multiple cancer types, including lung cancer, ovarian cancer, and thyroid cancer [Taylor et al., 2016]. Substantial fractions of these cancer types are grouped in the single cluster V (for which *BRAF* exhibits three interactions in Figure 5). The similarity of mutational profiles to other cancer types in cluster V suggests that they may be responsive to the same targeted treatment, such as *BRAF* inhibitors, provided the targeted gene or pathway is mutated.

## Discussion

We modelled mutational profiles with Bayesian networks, which capture the interactions between mutations, in a pancancer setting across 22 cancer types. Clustering of pancancer data can be highly insightful into the molecular similarities across cancer types [Ciriello et al., 2013, Hoadley et al., 2014]. In order to concurrently model cancer heterogeneity and mutational interactions we combined the Bayesian network approach with clustering in an integrated framework. The challenge for network clustering of large datasets resides in achieving reliable inference of networks with many nodes. Therefore we also developed novel methods for fast network learning for large data (Methods), so to extend the analysis to a couple of hundred important genes. Larger networks than those analysed may not lead to any further benefit, since many cancer mutations are quite rare and therefore unlikely to show strong interactions detectable in network modelling.

Mutations can be modelled at different levels, from the finer scale of individual mutation sites or hotspots [Chang et al., 2016] up to the pathway level. Higher resolution comes at the cost of lower frequencies, but this could be balanced out by combining aberrations by their molecular or pathway functions. A powerful alternative could be to use diffusion algorithms to condense aberrations to their affected subnetworks [Leiserson et al., 2015] or mediator genes [Dimitrakopoulos et al., 2018]. Here we focussed on mutation data summarised at the gene level, which contain a large amount of information. Mutational profiles are one genomic lens through which to identify molecular subtypes which can then complement other views based on, for example, copy number, expression and methylation profiles [Hoadley et al., 2014] and there may also be interactions across data modalities. Cancers can be highly heterogeneous, potentially harbouring distinct clones with different mutational profiles and prognostic signatures. Finer modelling of clonal structure and its impact on patient stratification present substantial challenges, as well as opportunities for more precise treatment options.

Furthermore we account for uncertainty in the network structure through a fully Bayesian approach (Methods). The networks learnt provide the key gene interactions characterising each cancer type (Figure 2) and a significantly extended view over just mutational frequencies, uncovering novel players and connections. Along with the edges, the networks also utilise the frequency of each mutation to quantify how similar or disparate the mutation patterns of patient samples are within and across cancer types (Figure 3).

Because the networks we infer are related to a class of clustering methods, we directly employed our models to re-cluster the TCGA data. We discovered and characterised 22 clusters in the data, distinct from the original cancer types but happening to match in number. New patient samples, including mutation profiles from different tumour types, can also be classified into the clusters, for which we provide a web application (Software). The new clusters significantly improve survival prediction, over and above that from the clinical and histopathological status of each patient sample. Integrating this information along with the mutational profiles during the clustering could further improve the survival prediction. Different types of mutations can occur and mutations also have different functional impact, which may affect their potential interactions, and may have different effects in different tissues. Accounting for such interactions could offer further refinements.

The Bayesian network modelling and clustering developed here provides us with novel insights into the mutation events and specifically their dependencies in cancer types and in novel cancer subgroups. These can be used as biomarkers which may be explored experimentally for therapeutic intervention.

## Software

A web interface to classify new patient samples is at https://cbg.bsse.ethz.ch/pancancer/ The package **BiDAG** for Bayesian network inference is at https://CRAN.R-project.org/package=BiDAG R code for the Bayesian network clustering and survival analysis is available at https://github.com/cbg-ethz/pancancer-clustering

## Author contributions

JK, GM, JB and NB designed the study. JK, GM and PS developed methodology. TT collated data. JK, TT and RG performed analyses. JK, TT, GM and RG drafted the article. GC and NB critically reviewed the manuscript. All authors approved the final version.

## Correspondence

Correspondence and requests for materials should be addressed to Jack Kuipers and Niko Beerenwinkel.

## Methods

### Cancer mutation data

Mutation annotation files were obtained through GDAC Firehose [Broad Institute TCGA Genome Data Analysis Center, 2016] for the 22 cancer types with sequenced primary tumour samples from more than 100 patients (see Supplementary Material Table S1), giving a total of 8,198 patient samples. The 16 most significantly mutated genes for each cancer type were collated to give a total of 201 genes considered. A binary matrix of the non-silent mutations in these genes across all patient samples was generated.

Putatively actionable targets are identified based on approved drugs or drugs currently undergoing clinical studies as collected by MyCancerGenome [Taylor et al., 2016], a manually curated precision cancer medicine knowledge resource. MyCancerGenome was queried through rDGIdb [Wagner et al., 2015, Thurnherr et al., 2016]. Functional annotation of genes was performed using the KEGGREST package [Tenenbaum, 2016].

### Graph inference

To quantify the extent to which a Bayesian network can explain a set of binary data, we use the BDe score [Heckerman and Geiger, 1995] where the state of a node *X* is determined by a different parameter *θ_Y_* for each configuration of its *m* parents **Y**: *P* (*X = 1* | ***Y***) = *θ_Y_* with a beta prior on each *θ_Y_* with hyperparameters 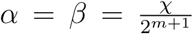 defined in terms of a single parameter χ which represents the number of pseudocounts added.

To make inference of the Bayesian network structure, which is a directed acyclic graph (DAG), we adopt a modified version of order MCMC [Friedman and Koller, 2003] as suggested by Kuipers and Moffa [2017]. For large networks we search over a reduced skeleton, initially found via the PC algorithm [Spirtes et al., 2000] and then expanded by MCMC search using the new software package **BiDAG** [Suter and Kuipers, 2017]. From the final skeleton, a sample from the posterior is obtained using partition MCMC [Kuipers and Moffa, 2017] optimised for such a skeleton [Suter and Kuipers, 2017].

### Prior edge knowledge

We obtained all human functional interactions from STRING [Szklarczyk et al., 2014], selecting those between the 201 genes included in this study (≈ 7000 interactions). Edges that were not in this STRING network were penalised by a factor 2 for the graph inference.

### Additional edge penalisation

When examining the networks learned from binary data, as in Figures 2 and 5 we additionally penalise all edges. Edge penalisation (by a factor of 2) has previously been examined with the marginal uniform prior [Scutari, 2016] and shown to improve network reconstruction. Based on simulation studies (Supplementary Material B) we find stronger regularisation further improves the accuracy for data mimicking the TCGA and employ a factor of 16 to regularise the network. We sample 100 DAGs from the posterior. Edges are only displayed if they appear in at least half of the posterior sample.

To select genes to display per cancer type or per cluster in Figures 2 and 5, for each gene we multiply its number of connections by its frequency, and choose the 20 genes with the largest product.

### Scoring samples against DAGs

For a given DAG the posterior distribution of each node *X* given its parent state ***Y*** is again a beta distribution with updated parameters 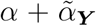 and 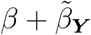, where 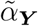 is the number of times *X* takes the value 1 when the parents are ***Y*** and 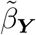 is the number of times it takes the value 0 in the data. Hence the likelihood

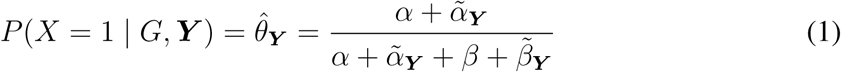
can be evaluated for each node of an arbitrary observed binary vector ***X***, providing a measure of fit of the observed vector to the DAG.

Given a sample of M DAGs from the posterior distribution *P*(*G* | *k*) of different cancer types (indexed by *k*), we can score each patient sample *D_i_* against each DAG *G_j_* (dropping the index for the cluster *k*). From the sample we can build the Monte Carlo approximation to the likelihood of the data for a given cluster *k*

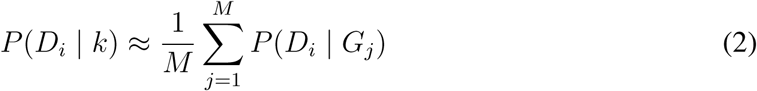
using the likelihoods in Equation (1). Under a uniform prior over cancer types, the likelihoods in Equation (2) are normalised to probability vectors over the collection of cancer types according to *P*(*k*|*D_i_*) ∝ *P*(*D_i_*|*k*)*P*(*k*). Similarities between the probability vectors of different patient samples are computed as their Jensen-Shannon divergence. Calculating this divergence between all pairs of patient samples provides a distance matrix between patient samples which we project into 2D with multidimensional scaling using the **cmdscale** command in **R**, as in Figure 3.

### Clustering with frequency information

We cluster the data with a mixture model. Given *K* graphs and parameter sets (*G_k_*, *θ_k_*), we assume that each patient sample *D_i_* is generated from one of *K* models depending on the value of a latent variable *Z_i_*

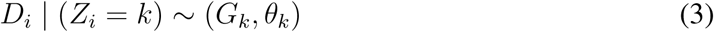
with

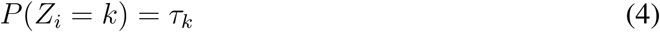

To start we allow no edges between the nodes. The different parameters *θ_k_* we learn for each empty graph *G_k_* only depend on the mutation frequencies in each group and do not account for any correlations between them. A Dirichlet prior is assumed on the cluster probabilities *τ* with all the parameters set to the value *χ*.

We find the maximum a posteriori (MAP) estimates of the mixture model parameters and the cluster membership probabilities of patient samples using the EM approach. This model-based clustering can also be seen as a latent class analysis [Dean and Raftery, 2010] where we include prior information. Without the prior (or taking the limit *χ* → 0), and without edges in the graph and hence independence between mutations, this clustering reduces to a Bernoulli mixture model.

### EM MAP algorithm

To find the MAP estimates for the mixture model we repeat the following steps: (1) given the membership probabilities, update the posterior parameters, and (2) given the updated parameters, update the probabilities. This is repeated until the probabilities and parameters no longer change.

At each step *t* of the algorithm we start with the current membership probabilities 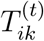 of each patient sample *i* being in cluster *k*. From the membership probabilities we can derive the cluster probabilities (as relative cluster sizes)

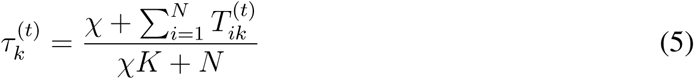
where *N* is the number of patient samples. A weighted version of Equation (1) gives the posterior means for each node j in the empty graph of each cluster:

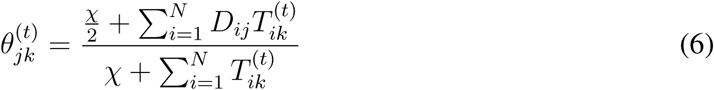
where *D_ij_*· is 1 if patient *i* exhibits mutation *j* and 0 otherwise so that it acts as an indicator function in the sum. From this we can directly evaluate 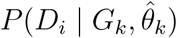

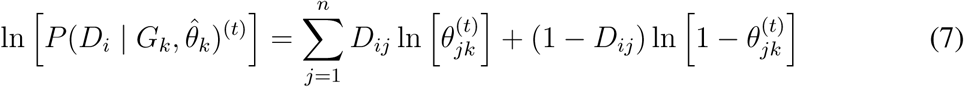
again using *D_ij_* as an indicator function.

Then we update the membership probabilities to start the next iteration

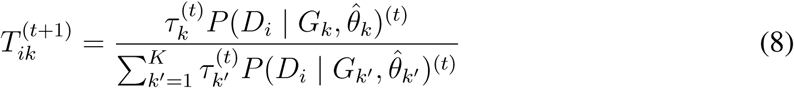
and repeat the iterations until the they no longer change.

The case where each patient sample has their membership probability equally spread across the clusters constitutes an unstable equilibrium for the mixture. To initialise the algorithm we therefore add a small random perturbation to this equilibrium to start with almost uniform probabilities 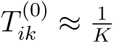.

### Cluster discovery

The global parameter χ quantifies the tradeoff between the within cluster variability and the number of clusters. Larger values of χ place less importance on each data point and lead to fewer larger clusters. Decreasing χ, the clusters break up leading to progressively more clusters. Since each clustering is stochastic, we run the clustering 240 times with different initial perturbations and retain the best. To find the optimal value of χ and number of clusters *K* we vary both and calculate a modified version of the AIC (Supplementary Material E). For each value of χ we choose the number of clusters with the lowest AIC and scan through a range of χ values. To select χ we chose the value whose clustering has the highest overlap (measured with the normalised mutual information) with clustering results for the range of χ.

### Clustering with structure learning

Allowing edges between the mutations in the DAG models refines the clustering. For each cluster we repeatedly learn the MAP DAG and reassign samples accordingly until convergence. In particular, we repeat the following steps: (1) given the current membership probabilities, learn the MAP graph structure, and (2) given the graph structure, repeat till convergence: (2i) given the membership probabilities, update the graph parameters, and (2ii) given the updated graph parameters, update the probabilities. We separate out learning the graph structure from updating its parameters since the latter is much less computationally demanding than the former. For given graph structures, the parameters and membership probabilities are continuous so the inner loop will converge. However since the graph space is discrete, periodic solutions for the outer loop are possible which can be alleviated by replacing the single graph in each cluster by a collection.

Once the clustering has converged and the final membership probabilities have been learnt, we sample 100 DAGs from the posterior of each cluster and visualise their connectivity and clustering features.

### Survival analysis

For each of the patient samples, we obtained clinical information through the TCGA web interface. To evaluate the power of the clusters derived from the mutational data for survival prediction, we employed the Cox proportional hazards model, with and without adjustment for age, stage and tissue type which are all significant predictors. The model fit with and without the cluster indicator was evaluated by the likelihood ratio test, based on the asymptotic *χ*^2^ distribution.

The possible values for stage were I, II, III, IV, ‘not assessable’ and ‘not available’. Performing a Cox regression with adjustment for age and tissue type, due to the low number of ‘not assessable’ samples keeping it as separate group from ‘not available’ did not lead to a significantly better predictions than their combination (likelihood ratio = 0.6; *p =* 0.3) and were hence combined into stage X. Combining stage I and II or stage III and IV led to significantly worse survival predictions and were hence kept separate. Therefore five levels (whose size is summarised in Supplementary Material Table S5) were retained for the stage covariate in the Cox regression.

### Overlap with functional network

To compare the STRING [Szklarczyk et al., 2014] network to the Bayesian networks inferred in this study we re-ran the analysis without using the functional STRING network as a prior. We then created the union of all edges in the Bayesian networks over all cancer types. We performed a permutation test on the overlap between these two networks by generating one million random permutations of the gene labels in the functional network. For each permuted network, we computed the mean overlap to derive the empirical *p*-value.

### Data availability

The mutational profiles and clinical information of the patient samples used in the study are available at https://github.com/cbg-ethz/pancancer-clustering, along with their cluster assignments.

### Code availability

The network inference code is available at https://CRAN.R-project.org/package=BiDAG.

The clustering and survival analysis code is available at https://github.com/cbg-ethz/pancancer-clustering.

